# Generation of a transgenic P*. cynomolgi* parasite expressing *P. vivax* circumsporozoite protein for testing pre-erythrocytic malaria vaccines in non-human primates

**DOI:** 10.1101/2024.12.26.630255

**Authors:** Maya Aleshnick, Shreeya Hegde, Charlie Jennison, Sebastian Mikolajczak, Ashley Vaughan, Derek Haumpy, Judith Straimer, Brandon Wilder

## Abstract

Malaria, caused by infection with *Plasmodium* parasites, exacts a heavy toll worldwide. There are two licensed vaccines for malaria as well as two monoclonal antibodies that have shown promising efficacy in field trials. Both vaccines and monoclonals target the major surface protein (circumsporozoite protein, CSP) of *Plasmodium falciparum*. Yet *Pf* is only one of the four major species of *Plasmodium* that infects humans. *Plasmodium vivax* is the second leading cause of malaria but *Pv* vaccine and monoclonal development lags far behind *P. falciparum* owing to the lack of basic preclinical tools such as *in vitro* culture or mouse models that replicate the key biological features of *P. vivax*. Notably among these features is the ability to form dormant liver stages (hypnozoites) that reactivate and drive the majority of *P. vivax* malaria burden.

*Plasmodium cynomolgi* is a simian parasite which is genotypically and phenotypically very close to *P. vivax*, can infect common research non-human primates and replicates many features of *Pv* including relapsing hypnozoites. Recently, a strain of *Pc* has been adapted to *in vitro* culture allowing parasite transgenesis. Here, we created a transgenic *P. cynomolgi* parasite in which the endogenous *Pc* CSP has been replaced with *Pv* CSP with the goal of enabling preclinical study of anti-*Pv* CSP interventions to protect against primary and relapse infections. We show that the *in vitro*-generated transgenic *Pc*[PvCSP] parasite expresses both serotypes of *Pv* CSP and retains full functionality *in vivo* including the ability to transmit to laboratory-reared *Anopheles* mosquitos and cause relapsing infection in rhesus macaques. To our knowledge, this is the first gene replacement in a relapsing *Plasmodium* species. This work can directly enable *in vivo* development of anti-*Pv* CSP interventions and provide a blueprint for the study of relapsing malaria through reverse genetics.

## Introduction

Malaria remains one of the most impactful infectious diseases worldwide, with approximately 263 million cases across 83 malaria-endemic countries in 2023 causing an estimated 597,000 deaths^1^. While the total number of malaria deaths fell precipitously from 2000 to 2015, this progress has plateaued in recent years^2^ highlighting the urgency of developing malaria interventions, including vaccines and therapeutics. Parasites in the genus *Plasmodium* are the causative agent of malaria, and infection of the vertebrate host is initiated when female *Anopheles* mosquitoes inject the sporozoite stage of the parasite into the skin while probing for a blood meal. These parasites travel to the liver, where each parasite infects a single hepatocyte and over ∼7-10 days forms tens of thousands of merozoites. These merozoites are released into the blood where iterative rounds of replication in red blood cells result in disease symptoms and transmission to the next vector (lifecycle reviewed in^3^). Several species of *Plasmodium* are responsible for the morbidity and mortality attributed to malaria, notably *P. falciparum (Pf)* and *P. vivax (Pv)*. While *Pf* is responsible for the majority of malaria-associated deaths worldwide^2^, *Pv* is the most widespread and exerts a significant toll in endemic areas, with nearly 5 million cases estimated in 2021^2^. In addition, *Pv* poses a unique challenge for elimination efforts due to the dormant hypnozoites that form in the liver which can reactivate months to years after the initial infectious mosquito bite^4^. These hypnozoites are a primary driver of disease and transmission as they are the source of ∼80% or more of *Pv* blood stage infections^5,6^.

While several points of the parasite lifecycle provide potential vaccine targets^7^, intervening at the pre-erythrocytic stage is an especially promising strategy as this stage is a bottleneck with relatively small numbers of extracellularly exposed parasites^8^. Abrogating parasite development at this stage would prevent all disease and transmission associated with the succeeding blood stage. Indeed, the only licensed malaria vaccines to-date, RTS,S/AS01 (Mosquirix^TM^) and R21/Matrix-M^TM^, both act at the pre-erythrocytic stage^9^. These vaccines both rely on antibodies against the circumsporozoite protein (CSP)—an abundant, essential protein with numerous critical functions during development in the mosquito to infection of the liver^10–16^. The general structure of CSP is conserved across *Plasmodium* species^15^, however the immunodominant “central repeat” region that is the target of neutralizing antibodies is variable between *Plasmodium* species with no serological overlap^17^. Thus, any CSP-based intervention must be developed specifically for each parasite species. In contrast to the two CSP-based *Pf* vaccines and the promising monoclonal antibodies CIS43LS and L9LS that are in clinical development^18–20^, there is a pronounced lag in the development of similar CSP-based interventions for *Pv*^21^.

The development of interventions against *Pv* has been hindered by the technical challenges of studying this parasite, namely the inability to culture the erythrocytic stages *in vitro* as is routinely done for *Pf*^22^. The result is that, aside from a few notable experimental infections of humans^23^, the study of *Pv* parasites at any stage must start by isolating blood from naturally infected persons. In addition, unlike the *Pf* CSP repeat region which is nearly invariant across field isolates, *Pv* has at least 2 major and one minor CSP repeat serotypes^24^ and so any CSP-based *Pv* vaccine will need to be at least bivalent to cover the majority of circulating parasites. These difficulties in developing a *Pv* model are exacerbated by lack of an *in vivo* model to study *Pv* liver stage biology, a better understanding of which will be critical for controlling and eradicating the disease.

While the murine *Plasmodium* species have allowed for substantial advances in *Plasmodium* research and the development of interventions, there is no mouse model of hypnozoites in which to study this critical parasite reservoir^22,25–27^. The logistical challenges in testing interventions in a controlled human malaria infection, as is commonplace for *Pf*^28^, leave gaps at every step of the development pipeline^29^. However, the simian malaria parasite *P. cynomolgi* is phylogenetically very closely related to *Pv*^30^ and mimics key aspects of *Pv* biology including a preference for reticulocytes at the blood stage^31^, the ability to form relapsing hypnozoites^31,32^, and infection of humans^33,34^. Importantly, *Pc* infects common laboratory non-human primates (NHPs) including rhesus macaques (RM), and the entire *Pc* life cycle can be completed between laboratory non-human primates and laboratory strains of Anopheles mosquitos—allowing consistent studies of infection and relapse. Thus, *Pc* has been a cornerstone of *Pv* research^25,31,32^. Indeed, hypnozoites were first described for *Pc*^35^ before their identification in *Pv* (reviewed in ^27^). This model was also critical in the identification of primaquine as the only pharmaceutical against liver-stage hypnozoites.^36^ The first vaccination study using a *Pc* sporozoite challenge showed this model to be stringent as a viral-vectored vaccine showed no efficacy^35^ and live-attenuated vaccination gave low but partial immunity^37^ reflecting what was seen with similar vaccines for *Pf* in humans. While the *Pc* animal model is ideal in many ways, the immunodominant repeat regions of *Pc* CSP and *Pv* CSP are divergent^38^ which precludes the direct assessment of *Pv* CSP-based interventions with an antibody component using *Pc* in NHPs. Recently, *Pc* blood stages have been culture-adapted^39^ which now allows for genetic manipulation without the need for propagation in NHPs as demonstrated by a recent study that introduced a drug resistance marker^40^. However, these *in vitro* parasites cannot yet be directly transmitted to mosquitoes as is done for *Pf*.

In this study, we aimed to overcome these practical limitations by creating a *Pc* parasite which expresses the *Pv* CSP protein and can complete the parasite life cycle between laboratory-reared *Anopheles* mosquitos and NHPs. This transgenic parasite fills a critical gap in the development pipeline of *Pv* pre-erythrocytic stage vaccines. Furthermore, our study provides the first example, to our knowledge, of gene replacement in *Pc* parasites that provides a blueprint for studies of relapsing Plasmodium parasites *in vivo* through reverse genetics.

## Methods

### Optimization of *in vitro* parasite culturing conditions

*Macaca mulatta* blood and serum was supplied by BioIVT (Westbury, NY) from US-born animals that had not been treated with antibiotics for at least a year. Samples were collected carefully in a sterile manner where needles were switched between extraction and transfer to vacutainer.

Whole blood was collected in heparin tubes (BD Biosciences: 367880) and blood for serum isolation was collected in serum tubes (BD Biosciences: 367815). Whole blood was spun down at 600g to remove the plasma and buffy coat, and red blood cells were washed thrice with RPMI 1640 with Glutamax, supplemented with 0.1% glucose, 30 mM HEPES and 200 µM hypoxanthine (*Pc* incomplete media) and stored at 4°C. A donor screen was carried out with 16 different NHPs to identify a serum lot supporting *Pc in vitro* culturing. Serum was collected, heat inactivated at 50°C for 1 hour with inversions every 15 minutes, filter sterilized, aliquoted and stored in −20°C. Donors were identified and booked for routine serum collection. The serum supporting the best *Pc* growth was selected for *in vitro* culture, and was added to *Pc* incomplete media at 20% v/v to create *Pc* complete media.

### In vitro culturing of *Plasmodium cynomolgi* Berok K2 parasites

Frozen vials of *Pc* Berok K2 parasites were received from NITD Singapore^39^. Cryopreserved parasites were thawed using a deglycerolization protocol consisting of three sodium chloride solutions (12%, 1.2%, and 0.9%) and maintained in *Pc* complete media in reduced oxygen (5% oxygen, 5% carbon dioxide and remaining nitrogen) incubators at 37°C. Regular media changes were performed and fresh *Macaca mulatta* red blood cells were supplemented in culture as needed to maintain 3-5% hematocrit.

### Parasite synchronization

*Pc* Berok K2 culture was centrifuged at 600g for 5 minutes to obtain a packed parasite culture. The pellet was incubated at room temperature for 20 minutes with 40x volume of sterile filtered 140 mM guanidine hydrochloride solution prepared in 20 mM HEPES buffer. The mixture was then washed by centrifuging at 600g for 5 minutes followed by washing twice with *Pc* incomplete media. The packed pellet was resuspended into media at 3-5% hematocrit and monitored for 48 hours until the majority of the parasites in the culture reached the schizont stage (within 40 to 48-hour window). To enrich for mature schizonts from the culture, density gradient purification was performed using Nycodenz as previously described^41^. In brief, the packed pellet was resuspended in 1mL *Pc* incomplete media to 50% hematocrit which was carefully layered over 5 mL of 55% Nycodenz solution diluted with phosphate buffered saline (PBS) in a 15 mL tube and centrifuged at 900g for 12 minutes without brake. A thin layer of schizont pellet was observed in the middle top layer and was gently extracted using a sterile Pasteur pipette. Schizonts were washed twice with *Pc* incomplete media and used for downstream transfections.

### Generation of a CRISPR/Cas9 plasmid for the editing of *P. cynomolgi*

The plasmid pYC-L2 was used as the backbone.^42,43^ The following components were amplified from *Pc* Berok K2 genomic DNA and ligated into the plasmid, replacing the *P. yoelii* components; EF1a bidirectional promoter, U6 promoter, HSP70 terminator, and replacement of the yFCU BiP 3 UTR with an mCherry BiP 3UTR fusion (Supplemental Table 1). The CSP cassette was ligated into this pCC-L2 plasmid.

### Generation of a chimeric *Pv* CSP cassette to replace *Pc* Berok CSP

To generate a plasmid targeting the CSP region of *Pc* Berok K2, a 3033 bp region was first amplified and sequenced, based on the available *Pc* M strain reference (PlasmoDB) (primer sequences in Supplemental Table 1). The chimeric *Pv* sequence was based on a previously published sequence^44^ with the plasmid kindly provided by Blandine Franke-Fayard and Chris Janse. An additional methionine after the signal peptide cleavage site in this sequence was removed with a fusion PCR to generate the final sequence encoding the 294aa *Pv* CSP chimeric repeat protein (**Fig 1A**, also see^44^).

**Figure 1:**
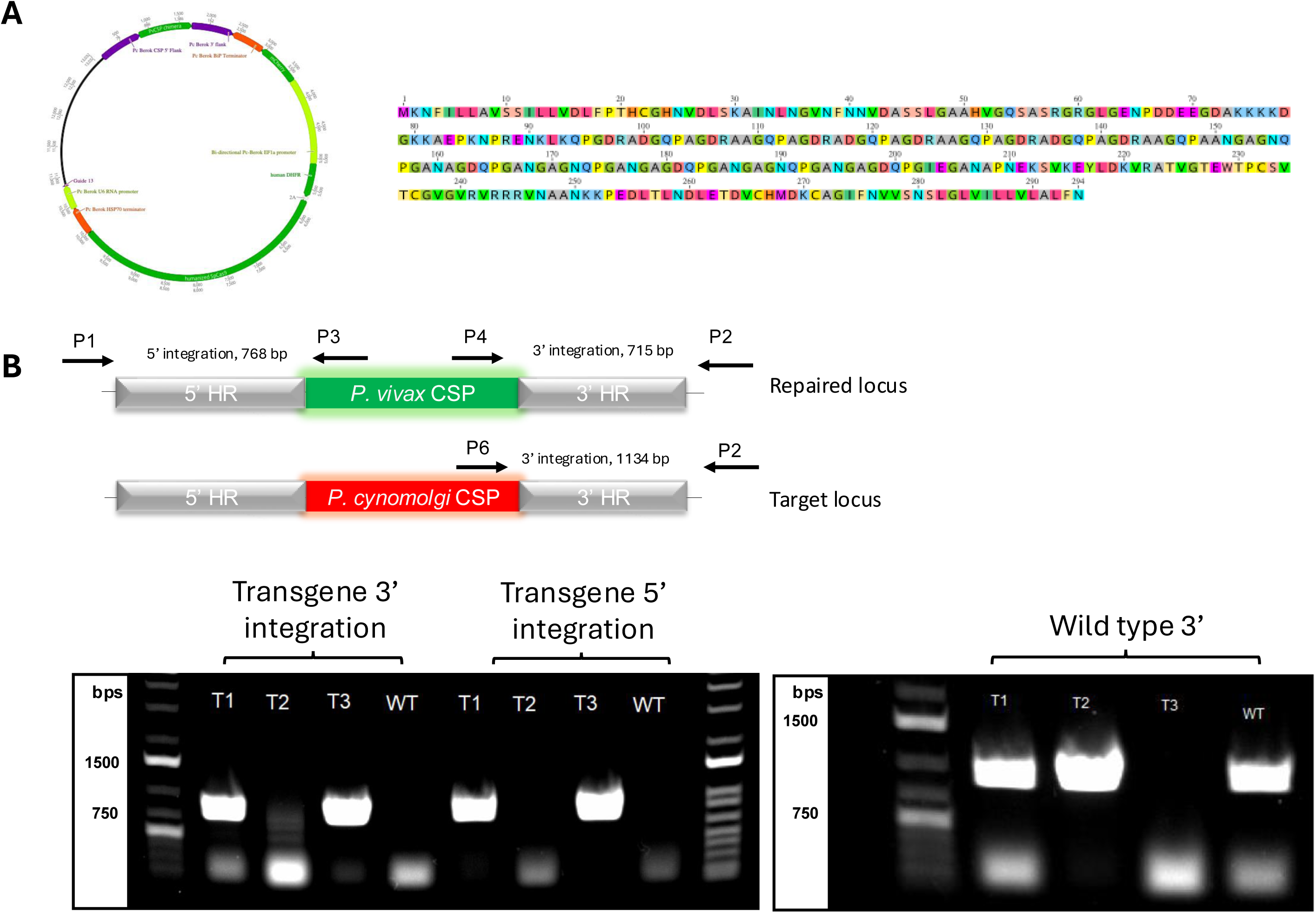
Construction and validation of a *P. vivax*-expressing *P. cynomolgi* parasite. A) Plasmid map and CSP sequence used for CRISPR *cas9* generation of the *Pc[Pv*CSP] transgenic parasite. B) PCR results confirming successful integration of the *Pv csp* gene replacing the native *Pc csp* after *in vitro* expansion of the transfected parasites. Top graphic indicates primers and expected amplicon size with genotyping primer pairs. Bottom images show PCR gels of transfected cultures T1-T3 with PCR for transgenic parasites on left and wild-type parasites on the right.

Based on the sequence data we had generated for the CSP region of *Pc* Berok, 700 and 685 bp regions of the 5’ and 3’ UTR, respectively, were amplified from genomic DNA and fused with the chimeric *Pv* CSP using primer overhangs in a fusion PCR. This CSP cassette was inserted into the pCC-L2 plasmid with SalI and NotI restriction sites (Supplemental Table 1). Using the online program CHOPCHOP (see ^45^), 3 guide sequences were selected and inserted into the pCC-L2-PvCSP plasmid downstream of the U6 promoter with the Esp3I restriction enzyme to generate pCC-PvChimera-CSP-G2/13/16 plasmids. Due to undesirable mutations identified in the CSP region of the G2 and G16 plasmids, plasmid G13 was used for transfections (named Pc_Pv plasmid, below).

### Nucleofection of *Pc* Berok K2 parasites

The protocol to transfect *Pc* schizonts was adapted from tools used to genetically modify *P. knowlesi*^46^. Nucleofections were performed using Lonza Amaxa 4D electroporator (Lonza) and P3 Primary cell 4D Nucleofector X Kit L (Lonza). A volume of 5-10 µL of purified *Pc* schizonts were incubated with 5 µL of 20 µg Pc_Pv CSP plasmid and 90 µL of P3 primary cell solution for 5 minutes. The mix was transferred to a 100 µL single nucleocuvette and immediately electroporated using program FP 138. Electroporated parasites were transferred to a 1.4 mL Eppendorf tube containing 500 μL prewarmed *Pc* complete medium supplemented with 25 μL fresh packed RBCs, and incubated at 37°C on a thermomixer, shaking at 650 rpm for 30 minutes. Transfected parasites were transferred to 24-well plate containing 1 mL *Pc* complete media and incubated as described above.

Media change was performed 24 hours after transfection and selection with 50 nM pyrimethamine was initiated; media was changed daily until the culture was free of parasites. Parasites were below limit of detection by Giemsa smears by day 7 of selection after which selection was stopped. *Pc* complete media was used for maintenance with media change done 3x weekly, 20% of total culture volume was cut and replaced with fresh blood and media weekly and periodic Giemsa smears were made to monitor culture health and detect parasites.

### Genotyping and sequencing of *P. cynomolgi* transgenic parasites

Sequencing of transgenic Pc[PvCSP] parasites from *in vitro* culture were set up using whole red blood cells from culture and thermo cycling conducted with Phusion Blood Direct kit (Thermo Fisher) to confirm recombinant locus. Sequencing of blood stage Pc[PvCSP] parasites from *in vivo* NHP infections were conducted using parasite pellet from *in vitro* cultures setup with NHP blood and genomic DNA isolated using the DNeasy blood and tissue kit from Qiagen, following manufacturer’s protocol. Three sets of primers were used for the PCR. The first set (P1-CATATCTGTACATGTCCATGTAGTGACC and P3-GAATACTACTCACGGCGAGC) were used to amplify a 786-bp fragment at the 5’ end of the recombinant locus Pc[PvCSP], and second set (P4-TATTCTGCTGGTGCTGGCCC and P2-ACGCAGTTTGCACACACCTGGC) were designed to amplify a 715-bp fragment at the 3’ end of the Pc[PvCSP] recombinant locus. The last set (P6-GTAAGAATGAGAAGAAAAGTTAGTGC and P2-ACGCAGTTTGCACACACCTGGC) were designed to amplify a 1134-bp fragment from the wild-type 3’ CSP locus. All three lines Pc[PvCSP]1, Pc[PvCSP] 2 and Pc[PvCSP] 3 along with *Pc* Berok K2 wild type parasites and plasmid DNA as controls were included in the PCR diagnostic.

### Mosquito infections

*Anopheles stephensi* mosquitoes were reared at OHSU or purchased from the University of Georgia SporoCore. Female mosquitoes were maintained in an incubator at 26°C, 80% humidity, and provided a sugar cube and water pad until the day of infection. Three to six-day old female mosquitoes were infected via a membrane feeder with blood drawn into a Lithium-Heparin vacutainer (Greiner Bio-One) from an infected RM. To prevent exflaggalation of the male gametes prior to uptake by mosquitoes, the blood was maintained at 39°C during transport from the non-human primate facility to the insectary. To minimize the potential contribution of anti-*Pc* antibodies, for some infections the RBCs were washed twice with RPMI and sera replaced with human AB+ sera. Mosquitoes were allowed to feed until all were either visibly engorged with blood (typically 75-90% of all mosquitoes) or lost interest. Infected mosquitoes were maintained at 26°C, 85% humidity and provided a sugar cube and water pad, changed daily. At 4-6 days post-infection, mosquitoes were provided a supplemental blood meal via an anesthetized mouse^47^. At 8-10 days post-infection, a sample of mosquitoes were dissected and midguts visualized under phase-contrast microscopy to quantify oocyst burden. At D16-19, mosquitoes were dissected and sporozoites quantified for downstream use, including infection of a naïve NHPs, immunofluorescence assays, and cryopreservation.

### Immunofluorescence assay (IFA) demonstration of sporozoite specificity for *P. cynomolgi* CSP antibodies

Freshly dissected sporozoites were enumerated, fixed in 4% paraformaldehyde for 20 minutes at room temperature, and following a PBS wash were loaded on a multiwell microscope slide at a 25,000 sporozoites per 30μL. Slides were allowed to air dry overnight, wrapped in aluminum foil, and stored at −80°C until permeabilization. After thawing, sporozoites were washed twice with 1× PBS and then permeabilized and blocked with 3% w/v bovine serum albumin plus 0.2% v/v Triton X-100 in 1× PBS for 1 hour at room temperature. Primary mouse monoclonal antibodies to *P. vivax* CSP (Pv-247-CDC and Pv-210-CDC antibodies were obtained through BEI Resources, NIAID, NIH: *Plasmodium vivax* Sporozoite ELISA Reagent Kit, MRA-1028K, contributed by Robert A. Wirtz) were diluted in 1× PBS/3% BSA and incubated overnight in a cold room in a humidified chamber. Following two washes with 1× PBS, fluorescent secondary antibodies were diluted in 1× PBS/3% BSA and incubated for 1 hour at room temperature in a humidified chamber and shielded from light. Nucleic acid was then stained with DAPI in 1× PBS for 10 min at room temperature. Sporozoites were washed three times with 1× PBS, then slides were mounted with one drop of ProLong Antifade reagent (ThermoFisher) and topped with a cover slip. After 30 minutes, slides were sealed with nail polish and stored at 4°C until imaging. Images were acquired using Olympus 1 × 70 Delta Vision deconvolution microscopy.

### Infection of non-human primates

Blood stage parasites preserved in glycerolyte were stored in liquid nitrogen until use and fresh sporozoites were used shortly after dissection. For infections initiated by blood stage parasites, a 200-500μL parasite stock originating from culture or previous RM infection was thawed and deglycerolization performed using saline washes as described above^48^. For infections initiated with sporozoites, 36,000 fresh sporozoites were washed once in sterile phosphate-buffered saline prior to infection. For all types of infections, the parasite pellet was suspended in 1mL sterile phosphate-buffered saline and injected intravenously via the saphenous or cephalic vein.

Beginning five days post-infection, a small blood sample was collected for detection of parasites via thin-smear microscopy. Complete blood counts were collected every 1-3 days during infection to monitor hematocrit and hematological factors associated with animal well-being. During peak infection, larger draws of several milliliters were collected to feed to mosquitoes and preserve parasites for validation of *Pv*CSP transgene integration and production of stocks for future infections. Animals were treated prior to a maximum parasitemia cutoff of 300,000 parasites/mL. To clear blood stage parasites, Coartem (20mg artemether/120mg lumefantrine, Novartis) was administered orally in a treatment course of 1 tablet twice daily for three consecutive days. For animals infected with sporozoites, semi-weekly monitoring was conducted to detect relapse infection from a reactivated hypnozoite. At the conclusion of the study, sporozoite-infected animals were treated with a single dose of Tafenoquine (150mg, GlaksoSmithKline, trade name Krintafel) to clear any remaining hypnozoites in addition to a course of Coartem.

All procedures involving animals were reviewed and approved by the Institutional Animal Care and Use Committee of the Oregon Health and Science University (IACUC protocol number: IP00002518).

### Quantification of parasites in NHP blood

Thin-smear microscopy to enumerate parasitemia was performed daily during patent infection and twice weekly during relapse monitoring to monitor the course of infection by Giemsa-staining and quantification of >20,000 parasites. To confirm the absence of parasites in the blood between blood-stage treatment and relapse, blood samples were preserved in RNALater™ solution (Invitrogen) and parasites quantified by qRT-PCR. In brief, extraction of nucleic acid from blood samples preserved in RNALater™ solution (200 μL in 400 μL, respectively) was performed using the MagMAX™-96 Total RNA Isolation Kit (Invitrogen). This was followed by pipetting of mastermix and template into 96-well plates for one-step amplification on the StepOnePlus™ instrument (Applied Biosystems). A Pan-Plasmodium 18S rRNA assay (Forward PanDDT1043F19: 5’-AAAGTTAAGGGAGTGAAGA-3’; Reverse PanDDT1197R22: 5’-AAGACTTTGATTTCTCATAAGG-3’; Probe: 5’-[CAL Fluor Orange 560]-ACCGTCGTAATCTTAACCATAAACTA[T(Black Hole Quencher-1)]GCCGACTAG-3’[Spacer C3]) alongside a housekeeping gene (TATA binding protein; Forward: 5’-GATAAGAGAGCCACGAACCAC-3’; Reverse: 5’-CAAGAACTTAGCTGGAAAACCC-3’; Probe: 5’-[FAM]-CACAGGAGC-[ZEN]-CAAGAGTGAAGAACAGT-[3IABkFQ]-3’) was used for quantification using the SensiFAST™Probe Lo-ROX One-Step Kit (Bioline). 0.2 μL of 10 μM probes and 0.8 μL of 20 μM primers were used for both targets. Cycling conditions were: reverse transcription (10 min) at 48°C, denaturation (2 min) at 95°C, and 40 PCR cycles of 95°C (5 sec) and 50°C (35 sec). Samples were run in triplicate.

## Results

### Generation of a *P. cynomolgi* parasite expressing the *P. vivax* circumsporozoite protein

A single plasmid carrying the *Pv* CSP chimera insert, *Cas*9 endonuclease and guide sequence was used to target the endogenous *Pc csp* locus (**Fig 1A**). Multiple transfection reactions were carried out and only 3 cultures (“T1-T3” in **Fig. 1B**) had parasites emerge 30 days post nucleofection. Culture T1 showed no presence of wild type parasite by PCR whereas Culture T2 had only wild type parasite. PCR of Culture T3 indicated a mixed population of wild type and transgenic parasites (**Fig 1B**). The selection marker, human dihydrofolate reductase, is not integrated in the locus but could be retained episomally. To confirm absence of episomal expression, Culture T1 was treated with 50 nM pyrimethamine. Parasite death was observed with culture clearance within 1 week indicating no presence of episomal plasmid. Parasites from this culture was used for all *in vivo* experiments. In summary, *Pc* Berok K2 wild type parasites were successfully modified to generate transgenic parasites expressing the *P. vivax* cirumsporozoite gene integrated in chromosomal DNA using CRISPR/Cas9 technology. This parasite is referred to as “*Pc*[*Pv*CSP]”.

### The *Pc*[*Pv*CSP] transgenic parasite completes the life cycle between rhesus macaques and *Anopheles* mosquitoes

We set out to determine if the *Pc*[*Pv*CSP] transgenic parasite infects macaques at the blood stage, can develop in the mosquito vector to form infectious sporozoites, and if these sporozoites are infectious *in vivo* with the ability to form relapsing hypnozoites (overview in **Fig 2A**). A cryopreserved stock of the *Pc*[*Pv*CSP] transgenic parasite produced *in vitro* was used to initiate an infection at the blood stage in RM1 for the purpose of parasite expansion and generation of cryopreserved blood stocks. After a prepatent period of 7 days, peak parasitemia was 37,927 parasites/μl (0.85%) at day 11, which began to decline at day 12 and continued to decline after treatment at day 13 according to planned protocol (i.e., not treated due to clinical treatment endpoint) (**Fig 2B**, leftmost panel). This NHP-cycled parasite was used to initiate an infection in RM2 to test the capacity of infected blood to infect mosquitoes and generate sporozoites. Parasitemia in RM2 was first identified on D8 post-infection, increasing to 77,905 parasites/μl until administration of antimalarial treatment on D12 (**Fig 2B**, second panel). This course of infection is similar to that seen in our lab for the *Pc* Berok strain WT parasite (data not shown). On days 11 and 12 post-infection, blood was fed to *A. stephensi* mosquitoes via a membrane feeder. A small number of mosquitos were dissected for detection of midgut oocysts at day 10 post-blood meal. Oocysts were not detected in any mosquitos dissected from the day 11 feed, and 6/10 mosquitos contained at least 1 oocyst in the mosquitos from the day 12 feed. Mosquito salivary glands from the Day 12 feed were dissected at day 16 post-blood meal which yielded 11,930 sporozoites/mosquito. We next intravenously injected 36,000 of these sporozoites into RM3 and monitored for primary and relapse infection (**Fig 2B**, third panel).

**Figure 2:**
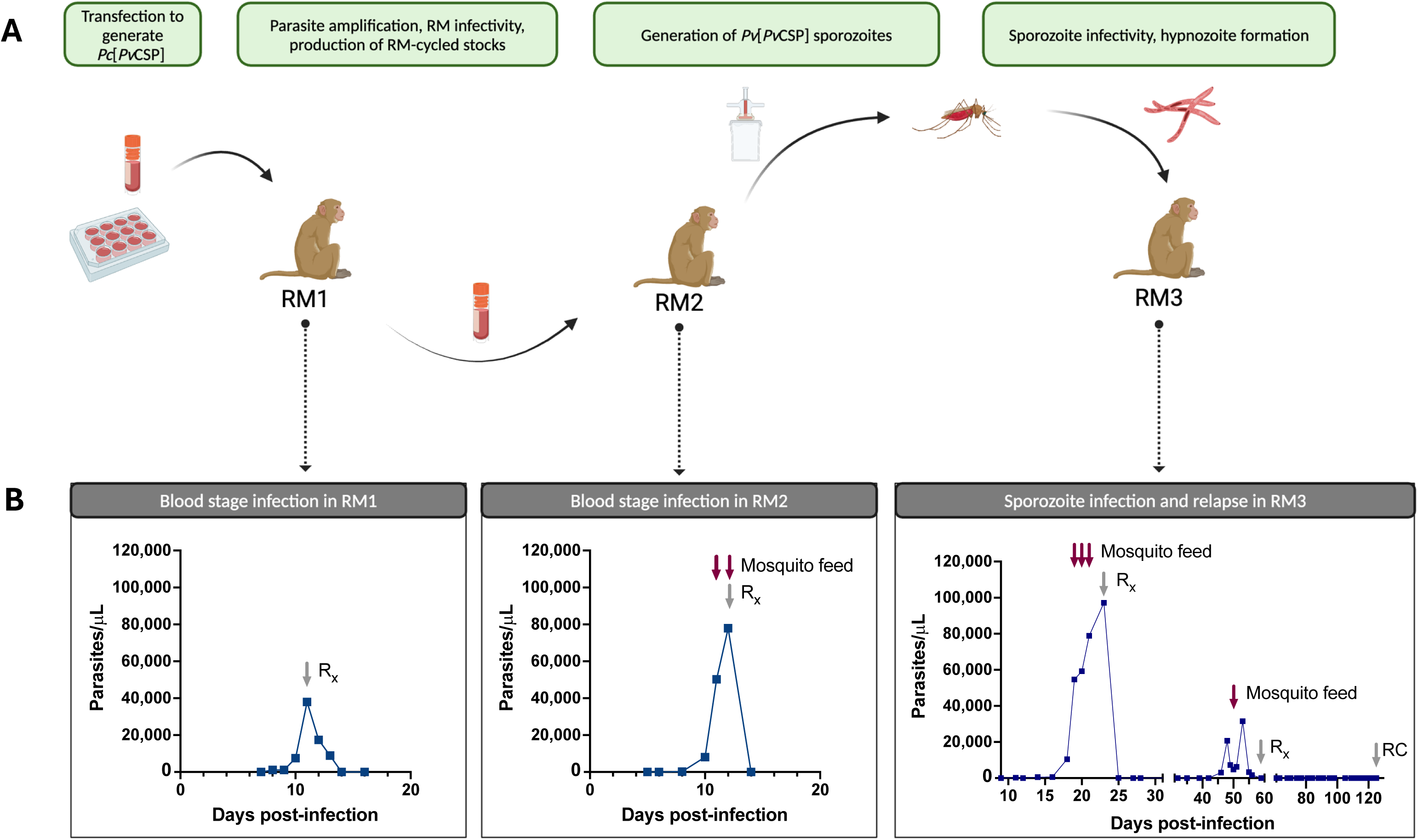
A *Pc*[*Pv*CSP] transgenic parasite completes the parasite life cycle between rhesus macaques and *An. stephensi* mosquitos including relapse. A) Overview of the NHP infections and mosquito feeds, with the purpose of each stage indicated above. B) Parasitemia curves demonstrating infectivity of the transgenic parasite at both blood and sporozoite stages. Parasitemia presented was quantified by thin-smear microscopy, parasite-negativity for animals infected with sporozoites (center panel) confirmed by qRT-PCR (see Supplemental Figure 1). *Pc*[*Pv*CSP] = *Plasmodium cynomolgi* parasite with endogenous circumsporozoite gene replaced with *P. vivax* sequence. RM = rhesus macaque. R_x_ = antimalarial treatment specific to clearing blood stage parasites (no action against hypnozoites). RC = radical cure with tafenoquine to clear hypnozoites.

Parasites were detected by microscopy starting on D11 post-infection and increased exponentially starting at D16. Mosquito feeds were conducted on days 19, 20, and 21 as the parasitemia approached 100,000 parasites/μl. Blood stage treatment with Coartem was administered on D23 and the animal became microscopy negative at D27. Importantly, the administration of Coartem does not eliminate hypnozoites, and so semi-weekly monitoring of parasitemia by thin-smear was initiated to monitor for hypnozoite reactivation after clearance of the primary infection. Parasites were again detected by microscopy on D46 (19 days after becoming microscopy negative post-Coartem), suggesting that the sporozoites maintained the ability to establish the relapsing infections. The relapse infection peaked at a lower parasitemia of 31,571 parasites/μl on D53 before slowly self-clearing, and Coartem was administered on D59 to ensure complete clearance of blood stage parasites. A single mosquito feed performed on the relapse infection yielded minimally-infected mosquitoes (702 sporozoites/mosquito) but confirm the ability of relapse parasites to infect the mosquito vector. On D125 post-infection, a radical cure treatment was administered to clear any potentially remaining hypnozoites.

Quantitative reverse-transcriptase polymerase chain reaction was used to provide a more sensitive measure of parasitemia, particularly between D28 and D39 to determine if we observed a true relapse rather than recrudescence of sub-microscopic parasite infection (**supplemental Fig 1**). Indeed, we found qRT-PCR to be more sensitive than microscopy with parasitemia persisting until D28 (vs. D27 via microscopy) but with a distinct return to baseline on D35. These results indicate a relapse, rather than recrudescence of a submicroscopic infection.

### Transgenic parasites maintain transgene integrity and protein expression *in vivo*

We next set out to ensure that we maintained transgene integration through the three RM using blood from the primary and relapse infections of RM3. Transgene integration was confirmed by polymerase chain reaction (PCR) using primers specific for the 3’ and 5’ ends of the *Pv* and *Pc* CSP proteins, on DNA isolated from parasite-enriched parental (*Pc* Berok K2) or RM-passaged transgenic (*Pc*[*Pv*CSP]) blood. The positive bands for the *Pc*[*Pv*CSP] parasite using *Pv*CSP primers, and lack of bands using parental *Pc* primers, indicates successful integration and complete replacement of the parental gene (**Fig 3A**). The transgenic *Pv*CSP contained the two serotypes, VK210 and VK247, that have been found across malaria-endemic areas and can co-exist in the same region and same patients ^39^. Thus, our transgenic parasite would have maximum utility only if it expressed both variants of CSP at the protein level to enable the preclinical assessment of anti-*Pv*CSP interventions. The expression of the chimeric *Pv*CSP protein was tested by immunofluorescent assay (IFA) using monoclonal antibodies (mAb) that binds to the VK210 (clone 2F2) and VK247 (clone 2E10) variants. In contrast to field isolates of sporozoites from Thailand and Peru that were only recognized by the anti-VK210 antibody, the *Pc*[*Pv*CSP] sporozoites were recognized by both monoclonals (**Fig 3B**). Together, these data indicate that the *Pc*[*Pv*CSP] parasite expresses both *Pv*CSP serotypes and maintains the *in vivo* capacity to complete the parasite life cycle, including relapse, in non-human primates. In addition to the first gene replacement of *Plasmodium* chromosomal DNA (vs. previous use of a maintained artificial chromosome^49,50^ or point mutation ^40^, this parasite will enable the first *in vivo* assessment of anti-PvCSP interventions to protect against primary and relapse infection in immune competent animals via a stringent parasite challenge.

**Figure 3:**
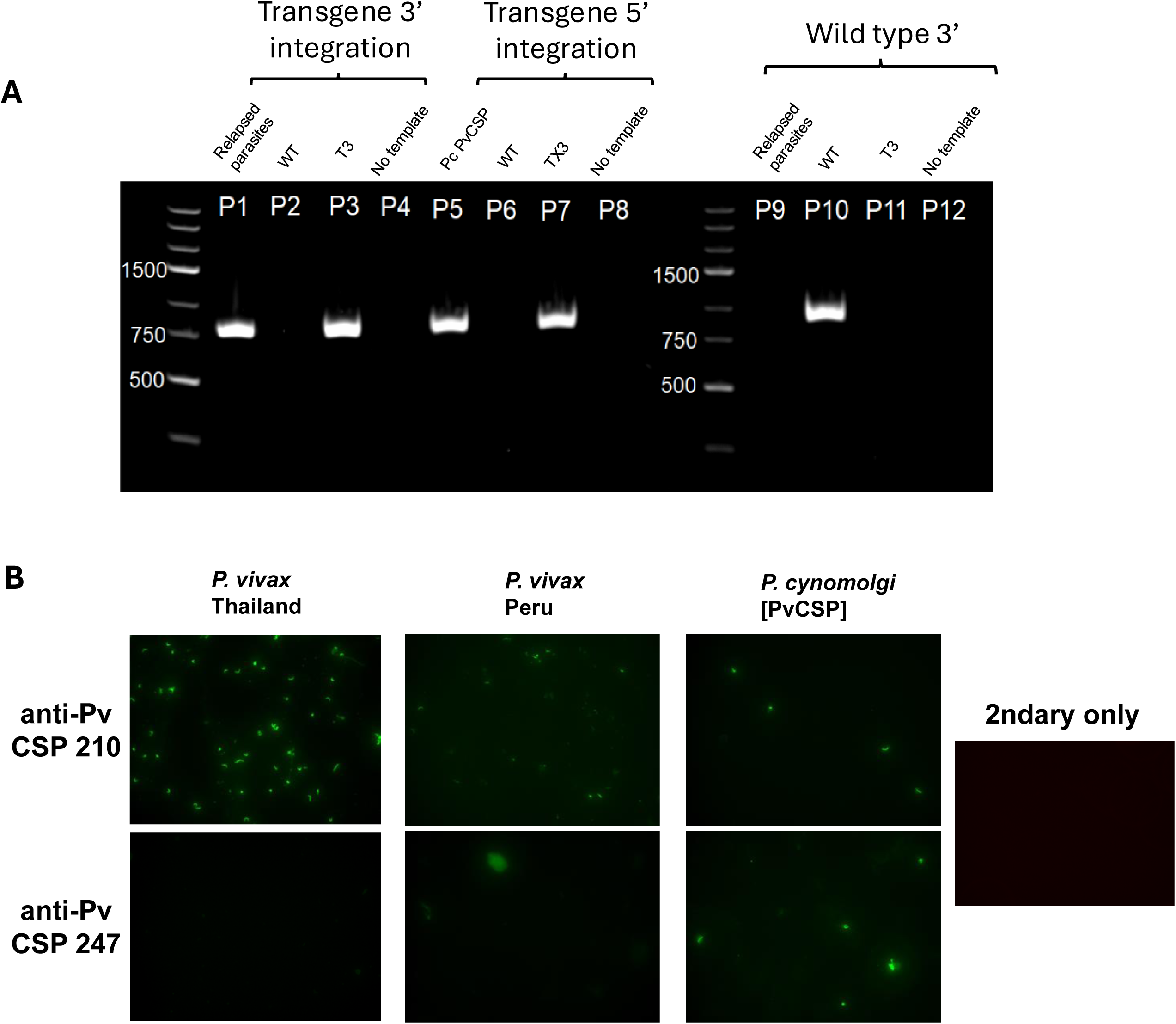
Confirmation of *Pv*CSP integration and protein expression following infection of NHPs and cycling in A*. stephensi* mosquitoes. A) PCR for integration of the *Pv csp* gene replacing the native *Pc csp* after cycling in three NHPs and *Anopheles* mosquitoes using infected blood samples from relapsed parasites, wild-type parental non-transgenic parasites (WT), Culture T3 transgenic parasites (T3), and no template sample using primer set described in Figure 1. B) Immunofluorescent imaging comparing expression of the two major *Pv*CSP variants in parasite field isolates from Thailand and Peru, both which predominantly express VK210, to our *Pc*[*Pv*CSP] transgenic parasite that expresses both VK210 and VK247. Secondary only control lacks any CSP-specific antibody.

## Discussion

In contrast to other *Plasmodium* species for which transgenesis has been commonplace, transgenesis of the closely related relapsing parasite *P. vivax* and *P. cynomolgi* has proven more difficult due to the lack of *in vitro* culture. This work capitalized on the recent *in vitro* adaptation of *Pc* which enabled CRISPR-mediated transgenesis without the need for parasite cycling and selection *in vivo*^39,40^. We used this opportunity to address a practical aim of a parasite that can be used to test CSP-based *P. vivax* interventions against primary and relapse infection. To this end, we have demonstrated the ability to perform parasite transgenesis and the capacity for our *Pv*CSP-expressing *P. cynomolgi* transgenic parasite to successfully complete the entire parasite lifecycle, including: infection of rhesus at both blood and liver stages, full development in the mosquito vector, and formation of hypnozoites that reactivate to generate relapse infections.

The circumsporozoite protein serves several critical functions in the mammalian host and in the mosquito vector^11^. While several CSP mutant parasites have been generated in *P. berghei*^51,52^, the necessity for correct function of this protein in sporozoite development is emphasized by an attempt to manipulate CSP in *P. falciparum* that resulted in parasites that abrogate at the oocyst stage^53^. Thus, the capacity of our transgenic parasite to not only complete development in the mosquito but also infect the NHP liver is encouraging even if unsurprising given the close phylogenetic relationship and overlapping hosts of *Pv* and *Pc*. To our knowledge, this is the first demonstration of gene replacement in a relapsing parasite and the only preclinical means of assessing the efficacy of anti-*P. vivax* CSP interventions in a relapsing model.

The steps in the formation and activation of hypnozoites has recently been identified as a key knowledge gap in understanding *P. vivax* biology, and thus in driving control and elimination^54^. This gap is perpetuated by the difficulty in obtaining *P. vivax* parasites which cannot be maintained in long-term culture. If *P. vivax* sporozoites are obtained, they can be used to infect cultured hepatocytes to observe dormant and replicating forms and have recently been used in humanized liver mice to demonstrate not only primary and relapse infection but susceptibility to monoclonal antibodies^55^. Still, the combination of logistical hurdles, difficulties in perpetuating the blood stage infection and absence of the immune system either *in vitro* or in humanized liver mice clouds our understanding of complex host-pathogen interactions. Previous studies have used the *P. cynomolgi* model to not only study the biology of infection, but to generate parasites carrying additional genes that allow for quantification and visualization *in vitro*^49^. As mentioned above, these studies utilized extra-chromosomal DNA to introduce new genes rather than the knockout, knock-in or mutation of genes possible with our CRISPR-based method. A more recent study used CRISPR-based mutation to knock-in a point mutation related to drug-resistance^40^. We hope our results are an important step to routine transgenesis of *Pc* parasites that enable a greater biological understanding of relapsing malaria.

In addition to a novel means to generate transgenic relapsing parasites, these studies will also advance vaccine and monoclonal antibody development for *P. vivax*. While valuable advancements in *P. vivax* vaccine testing have been made using transgenic murine parasites expressing the *P. vivax* CSP protein^56^, these rodent parasites don’t have the capacity to form hypnozoites and thus are not well-suited to test the effect of vaccine candidates on reducing the hypnozoite reservoir. Reactivation of hypnozoites is proposed to cause ∼80% of the observed *P. vivax* infections^57^. This means interventions that can reduce sporozoite infections of the liver, even if unable to completely abrogate infection, may have an outsized effect on *P. vivax* elimination. For example, while a vaccine or monoclonal antibody prophylactic capable of stopping 90-95% of *P. falciparum* sporozoites from reaching the liver still results in a blood stage infection 50% of the time^58^. Modeling the same reduction in liver burden for a *P. vivax* vaccine could reduce total blood stage infections by up to 90%^58^ and thus have an outsized impact on disease and transmission compared to *P. falciparum*. As mentioned above, a humanized-liver mouse model assessed the effect of passively transferred mAb to prevent primary and relapse infection ^26^, but this model is not suited to testing active immunization or interaction between antibodies and FcR-bearing cells as the humanized liver mice are severely immunocompromised. We hope our parasite can be used to test the effect of novel interventions in the context of a more complete representation of *Pv* biology.

While this transgenic parasite serves as an important step towards in the study of relapsing *Plasmodium* infections, there are several limitations. First, only the CSP gene of *P. cynomolgi* is replaced with the *P. vivax* homolog and thus it is only suitable for testing CSP-based interventions. There is growing consensus in the malaria vaccine community that a multi-antigen and multi-stage approach to vaccination will be required for long-term control or elimination with a vaccine or monoclonal antibody. As such, expanding the number of vaccine targets through repeated transgenesis or better defining the homology between the target proteins in *P. vivax* and *P. cynomolgi* as recently done with *Pv* AMA1^59^ will be critical.

Nonetheless, this study serves as an important proof-of-concept demonstrating that the *P. cynomolgi* parasite can be genetically manipulated to create chimeras expressing *P. vivax* proteins. In addition, while we demonstrate clearly that the *P. vivax* CSP protein is expressed on our *Pc*[*Pv*CSP] parasite and the *Pc*CSP gene was absent, we were unable to confirm at the protein level that the *P. cynomolgi* CSP was not co-expressed as we do not have a mAb which reacts against only *P. cynomolgi* CSP.

In summary, studying the liver stages of *P. vivax* is extremely challenging and these challenges drastically slow progress on *P. vivax* interventions. The expansion of animal models relevant for understanding *P. vivax* biology and testing interventions will be key for advancing interventions against *P. vivax*. This transgenic parasite provides a useful addition to the limited research tools available to study relapsing *Plasmodium* biology and to develop interventions against *P. vivax*.

## Acknowledgements

We thank the SporoCore at the University of Georgia, Atlanta, GA, for providing *Anopheles stephensi* mosquitoes to supplement those reared in-house.

Figure 2 was created in BioRender. Aleshnick, M. (2024) BioRender.com/x20k981

---

**Supplemental Figure 1:**
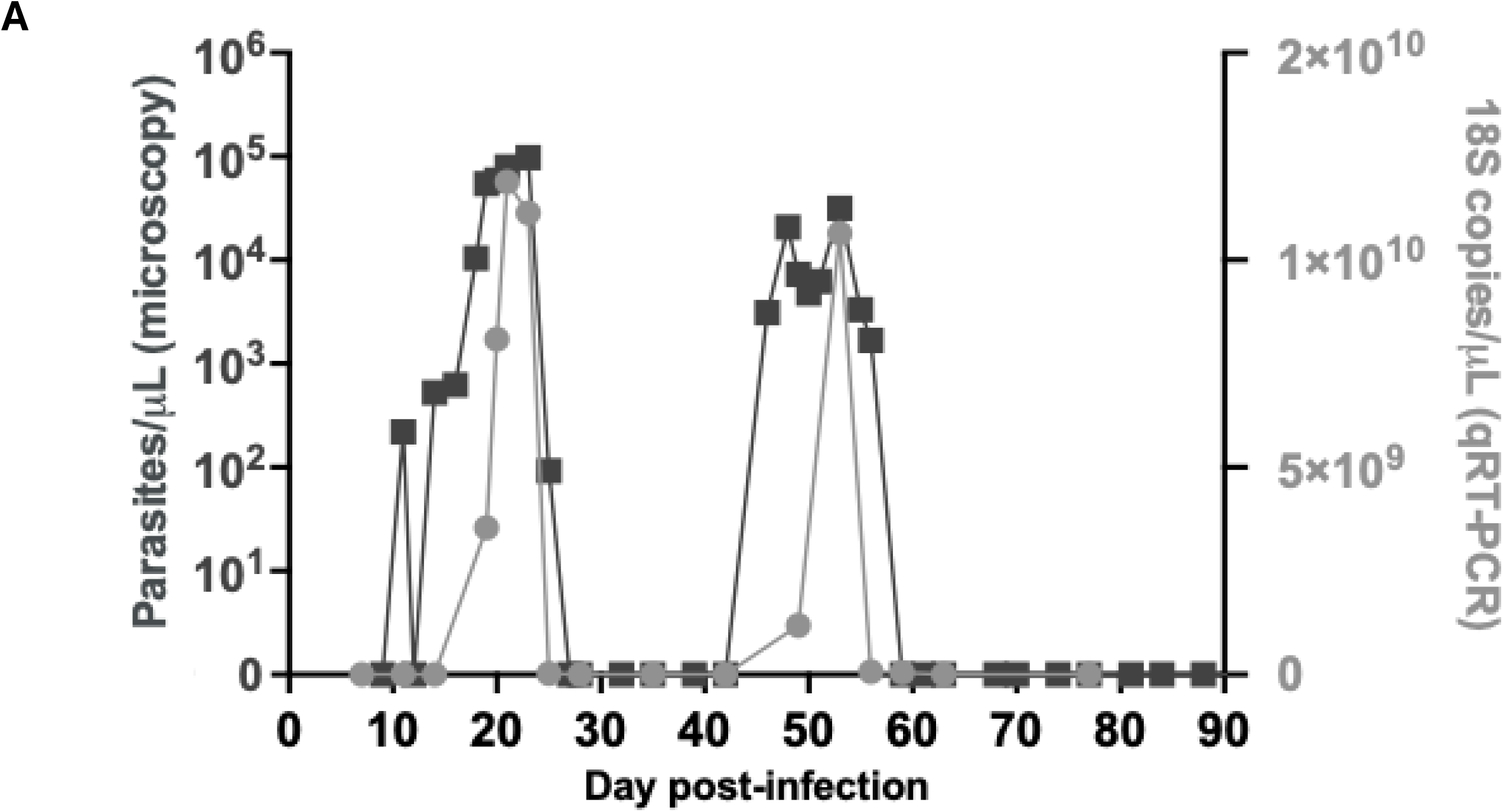
RM3 parasitemia by qRT-PCR and microscopy across primary and apse infections. Comparison of parasitemia as measured by microscopy (left axis) and T-PCR (right axis). PCR data points are the average of three triplicates.

**Supplemental Table 1:**
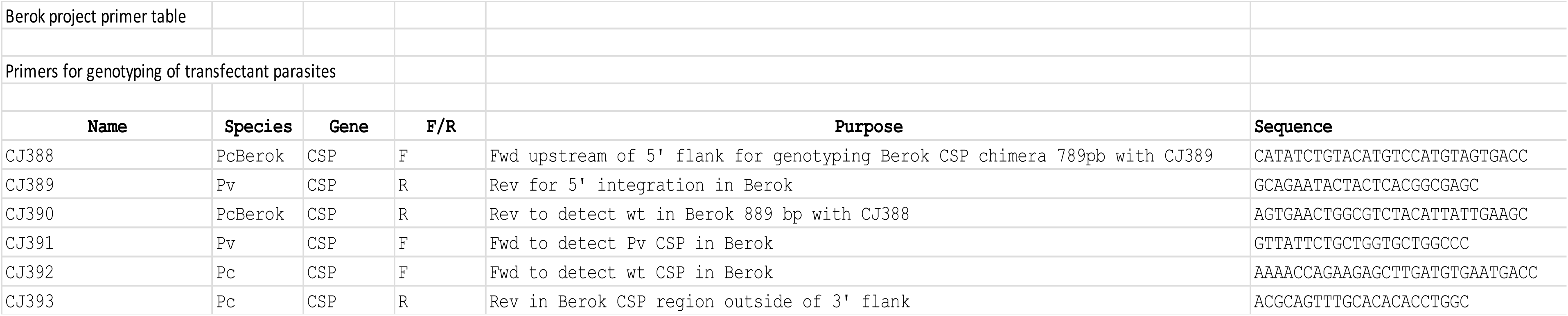
Primers used for genotyping parasites.

## References

1. World Malaria Report 2024. (World Health Organization, 2024).

2. World Health Organization. World Malaria Report 2022. https://www.who.int/teams/global-malaria-programme.

3. Simwela, N. V. & Waters, A. P. Current status of experimental models for the study of malaria. Parasitology vol. 149 729–750 Preprint at 10.1017/S0031182021002134 (2022).

4. Reyes-Sandoval, A. Plasmodium vivax pre-erythrocytic vaccines. Parasitol Int 84, 102411 (2021).

5. Adekunle, A. I. et al. Modeling the dynamics of Plasmodium vivax infection and hypnozoite reactivation in vivo. PLoS Negl Trop Dis 9, (2015).

6. Anwar, M. N., Hickson, R. I., Mehra, S., McCaw, J. M. & Flegg, J. A. A Multiscale Mathematical Model of Plasmodium Vivax Transmission. Bull Math Biol 84, 1–24 (2022).

7. Duffy, P. E. & Patrick Gorres, J. Malaria vaccines since 2000: progress, priorities, products. npj Vaccines vol. 5 Preprint at 10.1038/s41541-020-0196-3 (2020).

8. Smith, R. C., Vega-Rodríguez, J. & Jacobs-Lorena, M. The Plasmodium bottleneck: Malaria parasite losses in the mosquito vector. Mem Inst Oswaldo Cruz 109, 644–661 (2014).

9. Duffy, P. E. Current approaches to malaria vaccines. Curr Opin Microbiol 70, 102227 (2022).

10. Cohen, J., Nussenzweig, V., Vekemans, J. & Leach, A. From the circumsporozoite protein to the RTS,S/AS candidate vaccine. Hum Vaccin 6, 90–96 (2010).

11. Ménard, R. et al. Circumsporozoite protein is required for development of malaria sporozoites in mosquitoes. Nature 385, 336–339 (1997).

12. Zhu, F. et al. Malaria oocysts require circumsporozoite protein to evade mosquito immunity. Nat Commun 13, (2022).

13. Hopp, C. S. et al. Longitudinal analysis of Plasmodium sporozoite motility in the dermis reveals component of blood vessel recognition. Elife 4, (2015).

14. Stewart, M. J. & Vanderberg, J. P. Malaria Sporozoites Release Circumsporozoite Protein from Their Apical End and Translocate It along Their Surface. J Protozool 38, 411– 421 (1991).

15. Zhao, J., Bhanot, P., Hu, J. & Wang, Q. A Comprehensive Analysis of Plasmodium Circumsporozoite Protein Binding to Hepatocytes. (2016) doi:10.1371/journal.pone.0161607.

16. Arun Kumar, K., et al. The circumsporozoite protein is an immunodominant protective antigen in irradiated sporozoites. Nature 2006 444:7121 444, 937–940 (2006).

17. Davies, H. M., Nofal, S. D., McLaughlin, E. J. & Osborne, A. R. Repetitive sequences in malaria parasite proteins. FEMS Microbiol Rev 41, 923 (2017).

18. Kisalu, N. K. et al. A human monoclonal antibody prevents malaria infection by targeting a new site of vulnerability on the parasite. Nature Medicine 2018 24:4 24, 408–416 (2018).

19. Kisalu, N. K., et al. Enhancing durability of CIS43 monoclonal antibody by Fc mutation or AAV delivery for malaria prevention. JCI Insight 6, (2021).

20. Kayentao, K. et al. Subcutaneous Administration of a Monoclonal Antibody to Prevent Malaria. New England Journal of Medicine 390, 1549–1559 (2024).

21. Veiga, G. T. S. da, Moriggi, M. R., Vettorazzi, J. F., Müller-Santos, M. & Albrecht, L. Plasmodium vivax vaccine: What is the best way to go? Front Immunol 13, (2022).

22. Roobsoong, W., Yadava, A., Draper, S. J., Minassian, A. M. & Sattabongkot, J. The challenges of Plasmodium vivax human malaria infection models for vaccine development. Front Immunol 13, (2022).

23. Collins, K. A. et al. A Plasmodium vivax experimental human infection model for evaluating efficacy of interventions. J Clin Invest 130, 2920–2927 (2020).

24. González, J. M., Hurtado, S., Arévalo-Herrera, M. & Herrera, S. Variants of the Plasmodium vivax circumsporozoite protein (VK210 and VK247) in Colombian isolates. Mem Inst Oswaldo Cruz 96, 709–712 (2001).

25. Langhorne, J. et al. The relevance of non-human primate and rodent malaria models for humans. Malar J 10, 1–4 (2011).

26. Schäfer, C. et al. Partial protection against P. vivax infection diminishes hypnozoite burden and blood-stage relapses. Cell Host Microbe 29, 752–756.e4 (2021).

27. Voorberg-van der Wel, A., Kocken, C. H. M. & Zeeman, A. M. Modeling Relapsing Malaria: Emerging Technologies to Study Parasite-Host Interactions in the Liver. Front Cell Infect Microbiol 10, (2020).

28. Stanisic, D. I., McCarthy, J. S. & Good, M. F. Controlled Human Malaria Infection: Applications, Advances, and Challenges. Infect Immun 86, (2018).

29. Payne, R. O., Griffin, P. M., McCarthy, J. S. & Draper, S. J. Plasmodium vivax Controlled Human Malaria Infection – Progress and Prospects. Trends Parasitol 33, 141 (2017).

30. Galen, S. C. et al. The polyphyly of Plasmodium: comprehensive phylogenetic analyses of the malaria parasites (order Haemosporida) reveal widespread taxonomic conflict. R Soc Open Sci 5, (2018).

31. Simwela, N. V. & Waters, A. P. Current status of experimental models for the study of malaria. Parasitology 149, 729 (2022).

32. Galinski, M. R. Systems biology of malaria explored with nonhuman primates. Malar J 21, 177 (2022).

33. Muh, F. et al. Plasmodium cynomolgi: What Should We Know? Microorganisms vol. 12 Preprint at 10.3390/microorganisms12081607 (2024).

34. Collins, W. E., Coatney, G. R., Warren, M. & Contacos, P. The Primate malarias [original book published 1971]. (2003) doi:10.1126/science.177.4043.50.

35. Krotoski, W. A. et al. Relapses in primate malaria: discovery of two populations of exoerythrocytic stages. Preliminary note. Br Med J 280, 153 (1980).

36. Bykersma, A. The New Zoonotic Malaria: Plasmodium cynomolgi. Tropical Medicine and Infectious Disease 2021, Vol. 6, Page 46 6, 46 (2021).

37. Yongvanitchit, K. et al. Superior protection in a relapsing Plasmodium cynomolgi rhesus macaque model by a chemoprophylaxis with sporozoite immunization regimen with atovaquone-proguanil followed by primaquine. Malar J 23, (2024).

38. Verra, F. & Hughes, A. L. Biased Amino Acid Composition in Repeat Regions of Plasmodium Antigens. Mol. Biol. Evol 16, 627–633 (1999).

39. Chua, A. C. Y. et al. Robust continuous in vitro culture of the Plasmodium cynomolgi erythrocytic stages. Nat Commun 10, (2019).

40. Ward, K. E. et al. Integrative Genetic Manipulation of Plasmodium cynomolgi Reveals Multidrug Resistance-1 Y976F Associated with Increased in Vitro Susceptibility to Mefloquine. Journal of Infectious Diseases 227, 1121–1126 (2023).

41. Mohring, F., Hart, M. N., Patel, A., Baker, D. A. & Moon, R. W. CRISPR-Cas9 Genome Editing of Plasmodium knowlesi. Bio Protoc 10, (2020).

42. Zhang, C. et al. Efficient editing of malaria parasite genome using the CRISPR/Cas9 system. mBio 5, (2014).

43. Jennison, C. et al. Plasmodium GPI-anchored micronemal antigen is essential for parasite transmission through the mosquito host. Mol Microbiol 121, 394–412 (2024).

44. Miyazaki, Y. et al. Generation of a Genetically Modified Chimeric Plasmodium falciparum Parasite Expressing Plasmodium vivax Circumsporozoite Protein for Malaria Vaccine Development. Front Cell Infect Microbiol 10, (2020).

45. Labun, K. et al. CHOPCHOP v3: expanding the CRISPR web toolbox beyond genome editing. Nucleic Acids Res 47, W171–W174 (2019).

46. Mohring, F. et al. Rapid and iterative genome editing in the malaria parasite Plasmodium knowlesi provides new tools for P. vivax research. Elife 8, (2019).

47. Habtewold, T. et al. Plasmodium oocysts respond with dormancy to crowding and nutritional stress. Sci Rep 11, 3090 (2021).

48. Stanisic, D. I. et al. Development of cultured Plasmodium falciparum blood-stage malaria cell banks for early phase in vivo clinical trial assessment of anti-malaria drugs and vaccines. Malar J 14, (2015).

49. Voorberg-van der Wel, A. M., et al. A dual fluorescent Plasmodium cynomolgi reporter line reveals in vitro malaria hypnozoite reactivation. Communications Biology 2020 3:1 3, 1–9 (2020).

50. Voorberg-van der Wel, A., et al. Transgenic fluorescent Plasmodium cynomolgi liver stages enable live imaging and purification of Malaria hypnozoite-forms. PLoS One 8, (2013).

51. Othman, A. S. et al. The use of transgenic parasites in malaria vaccine research. Expert Rev Vaccines 16, 685–697 (2017).

52. Flores-Garcia, Y. et al. The P. falciparum CSP repeat region contains three distinct epitopes required for protection by antibodies in vivo. PLoS Pathog 17, (2021).

53. Marin-Mogollon, C. et al. Chimeric Plasmodium falciparum parasites expressing Plasmodium vivax circumsporozoite protein fail to produce salivary gland sporozoites. Malar J 17, 1–16 (2018).

54. Bassat, Q. et al. Key Knowledge Gaps for Plasmodium vivax Control and Elimination. Am J Trop Med Hyg 95, 62 (2016).

55. Schäfer, C. et al. A Humanized Mouse Model for Plasmodium vivax to Test Interventions that Block Liver Stage to Blood Stage Transition and Blood Stage Infection. iScience 23, 101381 (2020).

56. Salman, A. M. et al. Rational development of a protective P. vivax vaccine evaluated with transgenic rodent parasite challenge models. Scientific Reports 2017 7:1 7, 1–19 (2017).

57. White, M. T. et al. Modelling the contribution of the hypnozoite reservoir to Plasmodium vivax transmission. Elife 3, 1–19 (2014).

58. White, M., Amino, R. & Mueller, I. Theoretical Implications of a Pre-Erythrocytic Plasmodium vivax Vaccine for Preventing Relapses. Trends Parasitol 33, 260–263 (2017).

59. Winnicki, A. C. et al. Potent AMA1-specific human monoclonal antibody against Plasmodium vivax Pre-erythrocytic and Blood Stages. Nat Commun 15, 10556 (2024).

